# Seasonal Variation in Bacterial Community Structures of Mangrove Sediments

**DOI:** 10.1101/2023.05.23.541861

**Authors:** Nan Wang, Lu Liu, Zixiao Guo, Shaohua Xu, Rufan Zhang, Cairong Zhong, Suhua Shi, Ziwen He

## Abstract

Climate change globally and sea level rise affect the mangrove ecosystem. The high diversity and temporal heterogeneity of the mangrove ecosystem will lead to a high diversity of sediment bacterial community structure and function. However, seasonal variations and potential assembly mechanisms of sediment bacterial communities in mangrove ecosystems remain to be discovered. We collected rhizosphere sediments and bulk sediments from *Kandelia obovata* and *Aegiceras corniculatum* at three locations covering Dongzhai Harbour in spring, summer, autumn, and winter, and sequenced 16S rRNAs. The results indicated that the alpha and beta diversity of bacterial communities in mangrove sediments differed significantly between seasons, and the bacterial communities in rhizosphere sediments had smaller seasonal changes and were more stable than those in bulk sediment bacterial communities. The seasonal changes in carbon, nitrogen content, and pH were the main influencing factors. The stochastic process dominated the assembly of bacterial communities in mangrove sediments. The assembly of bacterial communities varies between seasons. We found that the proportion of dispersal limitation was significantly negatively correlated with the carbon and nitrogen content in the sediment. Compared with bulk components, the dispersal limitation of bacterial communities in rhizosphere sediments accounted for a lower proportion of community construction, which might be caused by higher carbon and nitrogen content conditions in rhizosphere sediments. We found that beta diversity based on Bray-Curtis distance was significantly positively correlated with dispersal limitation, which explained why the beta diversity of bacterial communities in rhizosphere sediments was significantly lower than that of bulk components. This study increases the understanding of the responses of mangrove bacterial communities to seasonal change and may be beneficial for the protection of mangrove ecosystems in the face of climate change.

## 1 Introduction

As the most productive and complex ecosystems, mangrove forests play essential roles in biogeochemical cycles, carbon sequestration, supporting intertidal biodiversity and regulating climate("Ecological role and services of tropical mangrove ecosystems: a reassessment," 2014). Located in intertidal zone, the environmental conditions of the mangrove ecosystem are dynamic and changeable due to the continuous changes of various physicochemical parameters such as salinity, temperature, tide, pH and nutrient content(G, D, K, & P, 2013). Due to its changeable physicochemical properties in the intertidal zone, the mangrove ecosystem is different from terrestrial and marine ecosystems(Alongi, 2014; Kathiresan & Bingham, 2001; Ruth, C, & E, 2010). Mangrove sediments contain a variety of microorganisms and invertebrates, which are breeding grounds, habitats and feeding areas for fish, mollusks, crustaceans, algae, crustaceans and birds(Gina, Patricia, & Yoav, 2001).

As we know, the mangrove ecosystem has active nutrient element circulation and rich organic matter content, so its microbial biomass and productivity are significantly higher than other ecosystems(Alongi, 1988). The bacterial community is a significant section of the mangrove ecosystem, and the abundant carbon and other nutrients in mangrove forest sediments support their activities(M., Paul, & Frank, 1993). In return, bacterial communities assist the organic matter degradation and nutrient cycling of mangrove forests. Mangrove leaves and wood are made mainly of lignocellulose components that are degradable by microorganisms("Effect of exported mangrove litter on bacterial productivity and dissolved organic carbon fluxes in adjacent tropical nearshore sediments," 1989; Moran & Hodson, 1989). The activity of bacteria is responsible for major nutrient transformations within a mangrove ecosystem. Complex interactions among these bacteria maintain the nutritional status and ecological balance of mangroves(Holguin et al., 2006).

Abiotic and biotic factors play an important role in determining the structure of bacterial communities in rhizosphere sediment. It has been found that the diversity and structure of bacterial communities are influenced by sediment properties and mangrove plants(Gogoi et al., 2019). The physicochemical conditions in mangrove forests are highly dynamic, the salinity, pH, nutrient availability and other conditions change with tides and distance to the coast(Feller et al., 2010; Gina et al., 2001). Salinity plays an important role in environmental filtration of the bacterial community in the ecosystem, and the bacterial community in low salinity conditions has higher bacterial diversity(Song et al., 2022). Salinity and pH are the main environmental factors that directly or indirectly determine bacterial community composition and diversity. Microorganisms, especially bacteria and fungi, are sensitive to the content of hydrogen ions in the environment(Zhang Lu et al., 2017). In addition, sediment pH can affect the solubility of nutrients in sediment solutions and the availability of nutrients to plants(Hu, Wang, & Yu, 2004). Lower pH will affect the interactions between sediments and bacteria(Rousk et al., 2010). Bacterial diversity in sediments decreases with the decrease in pH, and the bacterial community diversity is the highest under neutral conditions(Jizhong et al., 2002). The rhizosphere bacterial community of mangrove plants was significantly correlated with the content of key elements in the sediment. The assimilated sulfate reduction potential and oxidation potential of rhizosphere sediments are higher than that of bulk sediments(Ping, Yaping, Tiantian, Guoqiang, & Jun, 2018), and the relative abundance of Desulfobacteraceae is positively correlated with the content of phosphorus in the sediments and negatively correlated with the content of nitrogen in the sediments(Yanying et al., 2017). Plants can affect the soil environment through root exudates and litter. A large number of studies have described the mechanism of rhizosphere recruits bulk soil microorganisms, and highlighted the significant differences in the diversity and composition of rhizosphere and bulk soil bacterial communities(D. Bulgarelli et al., 2012; Edwards et al., 2015; Xiao et al., 2017).

The changes in climate and environment are also important factors affecting bacterial community diversity. In addition, seasonal changes will lead to dramatic changes in the physicochemical properties of mangrove ecology, which will affect the biological composition of mangrove ecosystems(Sundaramanickam et al., 2016). In spring and summer, Li studied the abundance of anaerobic ammonium oxidizing bacterial communities in mangrove sediments at different depths. The study showed that the diversity was higher in summer, and pH, salinity, the contents of nitrite, nitrate and ammonium salts in sediments may be the key factors affecting the community structure(Li et al., 2018). Behera studied the temporal and spatial heterogeneity of bacterial community structure and function in the Indian intertidal mangrove sediment. He found that the seasonal changes in bacterial community structure and metabolic pattern were largely explained by the seasonal abundance of different bacterial communities and the bacterial community composition was largely influenced by sediment salinity and organic carbon content(Behera et al., 2019).

One of the most fundamental problems in an ecosystem is how to generate and maintain diversity. It is widely acknowledged that there are two different ecological processes to explain the change of microbial community, namely deterministic (niche-based) and stochastic (neutral) processes(Hubbell; Leibold & McPeek, 2006). The neutral theory emphasizes that the attenuation of community similarity over space is related to the limited diffusion, without considering environmental differences between sites(Zhou & Ning, 2017). On the contrary, Niche theory claims that community similarity decreases with environmental distance, regardless of geographical distance(Zhou & Ning, 2017). The relative importance of these two process types in governing community assembly has been widely investigated(Dumbrell, Nelson, Helgason, Dytham, & Fitter, 2010; Jizhong et al., 2014; Stegen, Lin, Konopka, & Fredrickson, 2012; J. Wang et al., 2013). Deterministic processes based on niche theory and stochastic processes based on neutral theory are closely related to changes in community diversity and biogeographical distribution patterns(Hanson, Fuhrman, Horner-Devine, & Martiny, 2012; Zhou et al., 2015). Mangroves located in the intertidal zone provide good conditions for studying the effects of complex conditions on microbial community construction. Previous studies have shown that geographical distance affects the construction of mangrove microbial communities at different sites in southeastern China and that the differences between communities were mainly due to dispersal limitation between sites. The stochastic processes played an important role in the construction of mangrove bacterial communities(J. Wang et al., 2013). However, Zhang and Pan investigated mangrove microbial community construction in southeastern China and found that deterministic processes could better explain community construction(C. J. Zhang et al., 2019; Z. F. Zhang, Pan, Pan, & Li, 2021). In addition, the assembly processes of mangrove sediment bacterial community were influenced by a variety of complex factors, such as geographic distance, physicochemical properties of the sediment, and plant species. However, the influence of these factors on the construction process and how these factors influence the construction process remains to be elucidated. At present, most studies on mangrove bacterial communities under seasonal changes focus on the changes in bacterial community diversity and the effects of physicochemical factors on bacterial communities. Few studies try to explain the significant differences between mangrove bacterial communities and the reasons for the changes in bacterial community diversity under seasonal changes from the perspective of community construction.

Our research focused on (1) the seasonal changes in bacterial community diversity and composition of mangrove sediment and its correlation with the changes of intertidal climatic conditions and sediment physicochemical factors; (2) the seasonal variation of mangrove bacterial community assembly mechanism and the influence of abiotic environmental factors on bacterial community construction. We used high-throughput 16S rRNA gene sequencing to capture the bacterial communities of rhizosphere sediment of *Kandelia obovata* and *Aegiceras corniculatum*, as well as bulk sediments without plant roots. The neutral community model (NCM) proposed by Sloan was used to quantify the importance of stochastic processes(Sloan et al., 2006). The iCAMP analysis based on null model was used to estimate the assembly processes of mangrove sediment bacterial community(Ning, Deng, Tiedje, & Zhou, 2019). Based on the above studies, these results provide a meaningful reference for exploring the seasonal changes of mangrove bacterial community diversity and construction, and provide an example of mangrove bacterial community for in-depth exploration of the effect of various physicochemical factors on the community assembly processes. Further understanding of the seasonal changes of bacterial communities in mangrove ecosystems will provide a theoretical basis for the exploitation of microbial resources and the effective conservation of mangrove ecosystems.

## 2 Methods

### 2.1 Description of sampling sites and collection of sediment samples

Dongzhai Harbor is a drowned valley bay and located on the northeast of Hainan Island (Figure 1). We sampled rhizosphere sediments from individual trees of two mangrove species (*A. corniculatum* and *K. obovata*) and bulk sediments at three sites covering Dongzhai Harbor: Sanjiang (E110°38′, N19°56′), Yanfeng (E110°35′, N19°58′), and Tashi (E110°33′, N20°06′). For each plant species in each location, we sampled three rhizosphere sediment samples from three randomly selected trees at least 5 m apart from each other. At each location, we also collected three bulk sediments that are free of plant roots. We sampled samples in four seasons: spring (April), summer (July), autumn (November), and winter (January). Finally, we collected 108 sediment samples. All the sampled trees were mature individuals. To sample rhizosphere sediments of a tree, we collected plant roots at a depth of 5–15 cm. Rhizosphere sediments tightly adhering to the surface of roots were shaken off and fibrous roots were cut off with an aseptic scissor(Xu et al., 2018). To collect one bulk sample, we collected sediments at a depth of 5–15 cm within a quadrat of diameter 2 m. At each geographic location, the three bulk quadrats are at least 5 m apart from each other. Notably, both in sampling rhizosphere sediments of a tree and in sampling a bulk sample, we collected sediments from three rounded areas of diameter 30 cm within the tree or the bulk quadrat. All the sediments sampled from the same tree or a bulk quadrat were pooled as one sample. The sediment samples were packaged in sealed polythene bags and stored on ice before being transferred to a laboratory. In the laboratory, each sample was divided into two subsamples: the first subsample was stored at −80°C before nucleic acid extraction and the second subsample was stored at 4°C before examination of the physicochemical condition.

**Figure 1.**
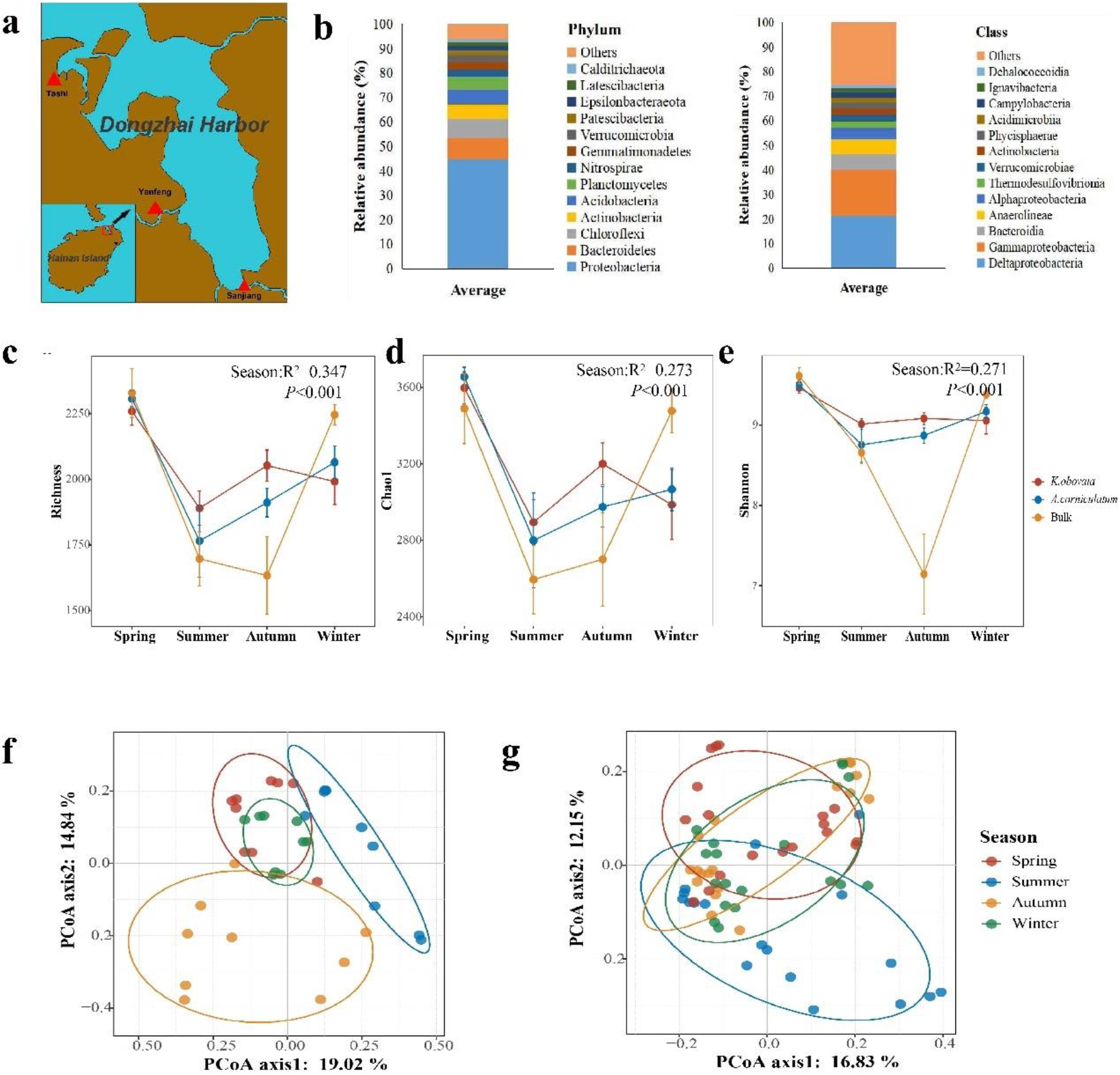
Bacterial community diversity and structure of three mangrove forests along Dongzhai Harbor in Hainan, China. (a) The map shows the sampled geographical locations in the Dongzhai National Mangrove Nature Reserve. (b) The average relative abundance of the bacteria in the mangrove sediment at phylum and class level. (d–e) The fluctuation of the microbial community richness index, Chao1 index and Shannon index in different seasons. (f) PCoA plots based on Bray-Curtis distance of bulk group. (g) PCoA plots based on Bray-Curtis distance of rhizosphere group.

### 2.2 Measurement of physicochemical conditions

The salinity of seawater was tested using an optical salinometer (Nanbei). The 1:2.5 sediment/water (dH_2_O) suspensions were shaken for 30 min before measuring the pH using a SevenEasy pH meter (Mettler Toledo). Similarly, the 1:5 sediment/water (dH_2_O) suspensions were shaken for 30 min before measuring the concentration of soluble salt. The salinity of sediment, which was determined by the Seven2Go S3 (Mettler Toledo), was determined to measure the amount of soluble salt. The contents of carbon, nitrogen, hydrogen, and sulfur were measured using a Vario EL cube V3.1.8 in CHNS mode (Elementar Analysensysteme GmbH) after the sediment was air-dried and filtered using a 200-mesh screen. We used 20–50 mg sediment of each sample for this measurement. The temperatures of the combustion furnace and the reduction furnace were set to 1150°C and 850°C, respectively. The CO_2_ and SO_2_ column desorption temperatures were set to 240°C and 220°C, respectively. The time duration of oxygenation was 120 s and the total test time for a sample was 10 min. The standard sample used for calibration was sulfanilic acid.

### 2.3 DNA extraction and 16S rRNA sequencing

For each sediment sample, the total genomic DNA was extracted from 0.5 g of sediment using a HiPure sediment DNA Kit B following the manufacturer’s protocol. PCR amplifications were conducted with the primer set 16S v34-F (CCTACGGGNGGCWGCAG) and 16S v34-R (GACTACHVGGGTATCTAATCC)(Deng et al., 2019; Ding et al., 2019), which amplified the V3–V4 region of the 16S rRNA gene. The PCR was performed in a 30 µl of reaction mixture according to the following process: initial denaturation at 94°C for 5 min, 25 cycles of denaturation at 94°C for 30 s, annealing at 57°C for 30 s, elongation at 68°C for 30 s, and final elongation at 68°C for 5 min. The amplicons were held at 4°C. All the obtained 16S rRNA amplicons were sequenced on an Illumina HiSeq platform.

### 2.4 Quality control of sequences and taxonomic classification

The 16S rRNA sequences were analyzed using VSEARCH2.7.1. The paired-end reads of the 16S V3–V4 region were merged using the FLASH program(Magoc & Salzberg, 2011). Quality control was performed to remove the following reads: (1) reads with quality scores less than 20; (2) reads with lengths less than 400 bp; (3) reads with ambiguous bases; and (4) singletons. The clean sequences were clustered into OTUs at a 97% similarity cutoff using the UPARSE clustering algorithm(Edgar, 2010). Chimeras were removed using the “gold” database (http://drive5.com/uchime/gold.fa). OTU sequences were aligned to the Greengenes database using PyNAST. Nonbacterial 16S sequences were discarded, and unaligned 16S sequences were discarded at a threshold of 75% identity. The taxonomic classification was performed at the 97% similarity level using the Ribosomal Database Project Classifier (version 2.11, http://rdp.cme.msu.edu/) against the SILVA v.132 16S rRNA database at a confidence of 80%(Q. Wang, Garrity, Tiedje, & Cole, 2007). The OTUs of the reads assigned to chloroplasts and archaea were removed. The Shannon, Chao1 and richness indexes were estimated to evaluate alpha diversity by subsampling 12255 reads (the smallest number of sequences with sufficient quality among all 108 samples).

### 2.5 Analyses of community diversity and structure

Community diversity was quantified using Shannon, Chao1 and richness indexes. Spearman’s correlation coefficients were calculated in the psych R library (version 2.2.5). PCoA was performed by the classical multidimensional scaling of beta diversity distance matrices using the *cmdscale* function in R 4.2.0. We performed the ANOSIM and PERMANOVA based on beta diversity distance matrices to test the dissimilarity of microbial communities among different samples. To compare the dissimilarity of Bray– Curtis distances and environmental distances in different rhizospheres, we performed the analysis of multivariate homogeneity of group dispersions (variances) using the betadisper implemented in the vegan R package (version 2.6.2). The environmental distances were calculated based on scaled Euclidean distances using the vegan R package (version 2.6.2). We used a Mantel test, which is based on Spearman’s correlation, to test the Bray–Curtis distances of bacterial communities are correlated with environmental distances. MRM analysis was performed using the vegan (v. 2.6.2), MuMIn (version 1.46.0), and ecodist (version 2.0.9) R packages.

### 2.6 Analyses of assembly mechanisms

We used the NCM to test the potential importance of neutral processes in community assembly, by comparing the detected frequencies of OTUs with their relative abundances in a wider metacommunity(Sloan et al., 2006). The nonlinear least-squares method was used in this prediction and the R^2^ value indicates the goodness of fit to the NCM(Sloan et al., 2006). Under this model, the estimated migration rate evaluates the probability that dispersal from a metacommunity replaces a random loss of an individual in a local community. Larger m values indicate that dispersal among microbial communities is less limited(Chen et al., 2019; Mo et al., 2021).

We also used the Phylogenetic-bin-based null model of the iCAMP R package (version 1.5.12) to infer the proportions of these community assembly mechanisms for individual phylogenetic bacterial taxa(Ning et al., 2020). We used the parameters d_s_ (phylogenetic signal threshold) =0.2 and *N*_min_ (minimum number) =24 in the binning step. In the Mantel test, the phylogenetic signal was considered significant if the Pearson correlation coefficient of the bins was R > 0.1 and *P* < 0.05. We conducted this computation using the “icamp.bins” function. The NRI was calculated using the “NRI.p” function. We first conducted iCAMP analysis by pooling samples from the four seasons for rhizosphere and bulk sediments, then we computed for rhizosphere and bulk sediments at each season.

## 3 Results

### 3.1 Seasonal distribution of environmental variables

In total, 108 sediment samples were collected from four different seasons in three sites (Figure 1a). The average values of environmental factors in each group are illustrated in Table S1. Among the 7 measured environmental variables, fast-changing environmental factors, such as sediment salinity (R^2^=0.617, *P*<0.001), pH (R^2^=0.163, *P*<0.001), and sediment moisture (R^2^=0.049, *P*<0.05), were largely influenced by the seasons. In summer, the sediment salinity was significantly higher, yet the pH and sediment moisture was significantly lower than in other seasons (Figure S1 a-c). In contrast, the total contents of carbon, hydrogen, nitrogen, and sulfur in sediments were relatively stable (Figure S1 d-e). Besides, the total contents of carbon and nitrogen in mangrove rhizosphere sediments were higher than the bulk (ANOVA, *P*<0.05).

### 3.2 Seasonal distribution of the sediment bacterial community composition

A total of 3,086,888 sequences of 16S rRNA were obtained across the 108 sediment samples. Overall, 10,491 operational taxonomic units (OTUs) were annotated at 97% identity.

For bacterial alpha diversity, season explained 34.7%, 27.3% and 27.1% of the Richness index, Chao1 and Shannon index variation in sediment bacteria. These three indexes were both significantly highest in spring, then decreased in summer, and finally recovered in winter (*P*<0.05, ANOVA) (Figure 1 c-e). Besides, the alpha diversities for rhizosphere bacterial communities of two mangrove species were consistent with seasonal fluctuations, and the fluctuation of the rhizosphere was smaller than that of bulk, which means, the rhizosphere bacterial community in mangrove sediment was more resilient to environmental disturbance than that in bulk. When considering alpha diversity between rhizosphere and bulk groups, the alpha diversity of rhizosphere samples was significantly higher than that of bulk samples (*P*<0.05, ANOVA).

In addition to the alpha diversity analysis, we examine the bacterial community composition according to seasons and species. At a 97% taxonomy identity threshold, most sequences of 16S rRNA were mainly assigned to Proteobacteria (44.71%), Bacteroidetes (8.66%), and Chloroflexi (7.90%) at the phylum level. At the class level, Deltaproteobacteria was dominant (belonging to Proteobacteria), followed by Gammaproteobacteria and Bacteroidia. The relative abundance of the order Desulfobacterales was highest, followed by Anaerolineales, Bacteroidales, Betaproteobacteriales, and Myxococcales (Figure 1b). When regardless of the seasonal factor, the relative abundance of Deltaproteobacteria was significantly lowest in spring, Planctomycetes, Patescibacteria, and Latescibacteria were highly abundant in spring, conversely (*P*<0.05). In addition, Verrucomicrobia were more abundant in spring and summer than that in autumn and winter, Actinobacteria did the reverse (*P*<0.05). Moreover, the relative abundance of Chloroflexi was significantly enriched in winter (*P*<0.05). It was noted that there were significant changes in the relative abundance of the top 13 bacterial phyla in mangrove sediment samples across different seasons (Figure S2). Furthermore, we also compared the composition of mangrove sediment bacterial communities in rhizosphere and bulk samples. Bacterial communities in the rhizosphere showed a significant increase in relative abundance for taxa belonging to Deltaproteobacteria, Patescibacteria, and Calditrichaeota (FDR < 0.05, Wilcoxon rank-sum test). However, higher relative abundances of Gammaproteobacteria and Actinobacteria were detected in the bulk sediment than those in the rhizosphere sediment (*P*<0.05, ANOVA).

### 3.3 Seasonal distribution of the beta diversity in sediment bacterial community

Using PERMANOVA and ANOSIM based on weighted UniFrac, unweighted UniFrac and Bray-Curtis distance matrices, bacterial communities were significantly distinguished by habitat (bulk sediment vs. rhizosphere sediment) (*P*<0.001). However, there was no significant difference between the rhizosphere bacterial communities of the two mangrove species (Table S2). So, we put two kinds of samples of *Kandelia obovata* and *Aegiceras corniculatum* together as one rhizosphere group. Based on the three distances, we performed PCoA analysis on the rhizosphere group and bulk groups respectively (Figure 1 f-g; Figure S3). For the bulk group, the results of PCoA based on weighted UniFrac, and Bray-Curtis distance showed that the bacterial communities of autumn were separated from other seasons. Besides, the bacterial abundance was related to the calculation of the above two distances. The abundance of the bulk group was significantly lowest in autumn, which could lead to this result. Notably, ANOSIM (permutations=999) based on weighted UniFrac, unweighted UniFrac and Bray-Curtis distance revealed that the difference of bulk bacterial communities between any two seasons was significant (*P*<0.05) (Table S3-4). In addition to the rhizosphere group, the result of ANOSIM based on Bray-Curtis distance was consistent with the above, and under the weighted UniFrac and unweighted UniFrac distance, the difference between any two seasons is also significant except for one group (ANOSIM, *P*<0.05) (Table S3-5). In general, the seasonal variability of bacterial beta diversity in mangrove sediment presents clearly.

### 3.4 Seasonal distribution of the sediment bacterial community structure

Moreover, we visualized the beta diversity among different groups. The beta diversity of the bacterial community (Bray-Curtis dissimilarity index) in the rhizosphere sediment was notably lower than that in the bulk sediment (Krusktal-Wallis test, *P*<0.001), and there are consistent results in different seasons, indicating the convergence in bacterial community composition of the rhizosphere sediments (Figure 4 a-b). Comparing Bray–Curtis dissimilarities of different seasons in bulk and rhizosphere bacterial communities, the results showed that the internal variation of the bulk group in different seasons was more apparent than that in the rhizosphere group, with obvious fluctuations (Figure 4 c-d). Overall, the beta diversity of bacterial communities across different seasons fluctuated in bulk sediment more dramatically than that in rhizosphere sediment.

Venn diagrams were performed to show the distribution of all detected OTUs in different seasons (Figure 4 e-f). The OTUs shared by four seasons in the rhizosphere and bulk group accounted for the total OTUs reached 50.83% and 40.96%, respectively. For most species, the changes in different seasons of their existence were relatively small. The result suggested that there was no significant difference in species pools between different seasons. The number of OTUs shared between seasons in bulk group was much fewer than that in the rhizosphere. In terms of the OTUs exclusive to four seasons, the result was opposite. According to the above results, we found that the species composition in bulk group had a bigger divergence than rhizosphere group, which also confirmed the previous conclusion that bulk bacterial communities showed higher beta diversity. In summary, considering beta diversity, bulk bacterial communities had significant differences across different seasons, and the rhizosphere group was relatively stable, suggesting that maybe the plant roots reduce the seasonal fluctuation.

### 3.5 Correlations between diversity of microbial community and environmental factors

The Mantel test based on the matrix of weighted UniFrac, unweighted UniFrac and Bray-Curtis distances indicated that the microbial community in mangrove sediment was significantly correlated with physicochemical conditions (r=0.4719; *P*<0.001, r=0.506; *P*<0.001, r=0.317; *P*<0.001). MRM analysis was performed to identify the key physicochemical factors which shape the bacterial communities. The result showed that the structure of bacterial community was significantly influenced by pH (coefficient=0.138, *P*=0.0004), moisture (coefficient=0.141, *P*=0.0001) and sulfur content (coefficient=0.102, *P*=0.0014). To further explore the correlation between key environmental factors and bacterial communities, linear regression analysis was carried out in this study. The figure S4 reflected that the Chao1 index and OTU richness showed a significant positive correlation with pH and moisture (*P*<0.05), but not significantly with sulfur content.

### 3.6 The construction process of mangrove bacterial community

The neutral community model (NCM) was used to evaluate the potential contribution of the stochastic process to the mangrove sediment bacterial community assembly, and the R2 values represent the goodness of fit to the NCM(Logares et al., 2013; Sloan et al., 2006). As shown in figure 3, the NCM showed 83.05 and 69.80% of explained bacterial community variance in rhizosphere group and bulk group, respectively, suggesting the stochastic process played important roles in the mangrove sediment system. Besides, the value of m was higher for bacterial taxa in rhizosphere (0.513) than that in bulk (0.359), indicating the extremely lower capability of microorganisms to disperse among bulk sediment. To further identify the process of rhizosphere and bulk bacterial community construction among different seasons, the phylogenetic-bin-based null model analysis (iCAMP) was performed. As shown in table S6, the stochastic process played a dominant role in the community construction of rhizosphere and bulk sediment bacterial communities, which is consistent with the results of the NCM. Furthermore, we identified the assembly processes of these bacterial communities by quantifying the importance of five assembly processes (i.e., homogeneous selection, heterogeneous selection, dispersal limitation, homogeneous dispersal and undominated processes). In both analyses conducted by sediment type and by seasons, the dispersal limitation and homogeneous selection have shown substantial contributions but the heterogeneous selection and homogenizing dispersal have contributed much less (Figure 3 c, Table S6). In spring and autumn, the contribution of dispersal limitation in the construction of bulk group was significantly higher than that of the rhizosphere group. This corresponds to the higher migration rates of rhizosphere group than bulk group in the spring and autumn found during the fitting of the NCM, indicating that rhizosphere bacterial communities have higher diffusion ability compared to bulk communities. There were significant differences in the process of community construction between different seasons (Figure 3c). Whether in the rhizosphere bacterial community or the bulk bacterial community, the homogeneous selection in summer is significantly lower than in the other three groups.

**Figure 2.**
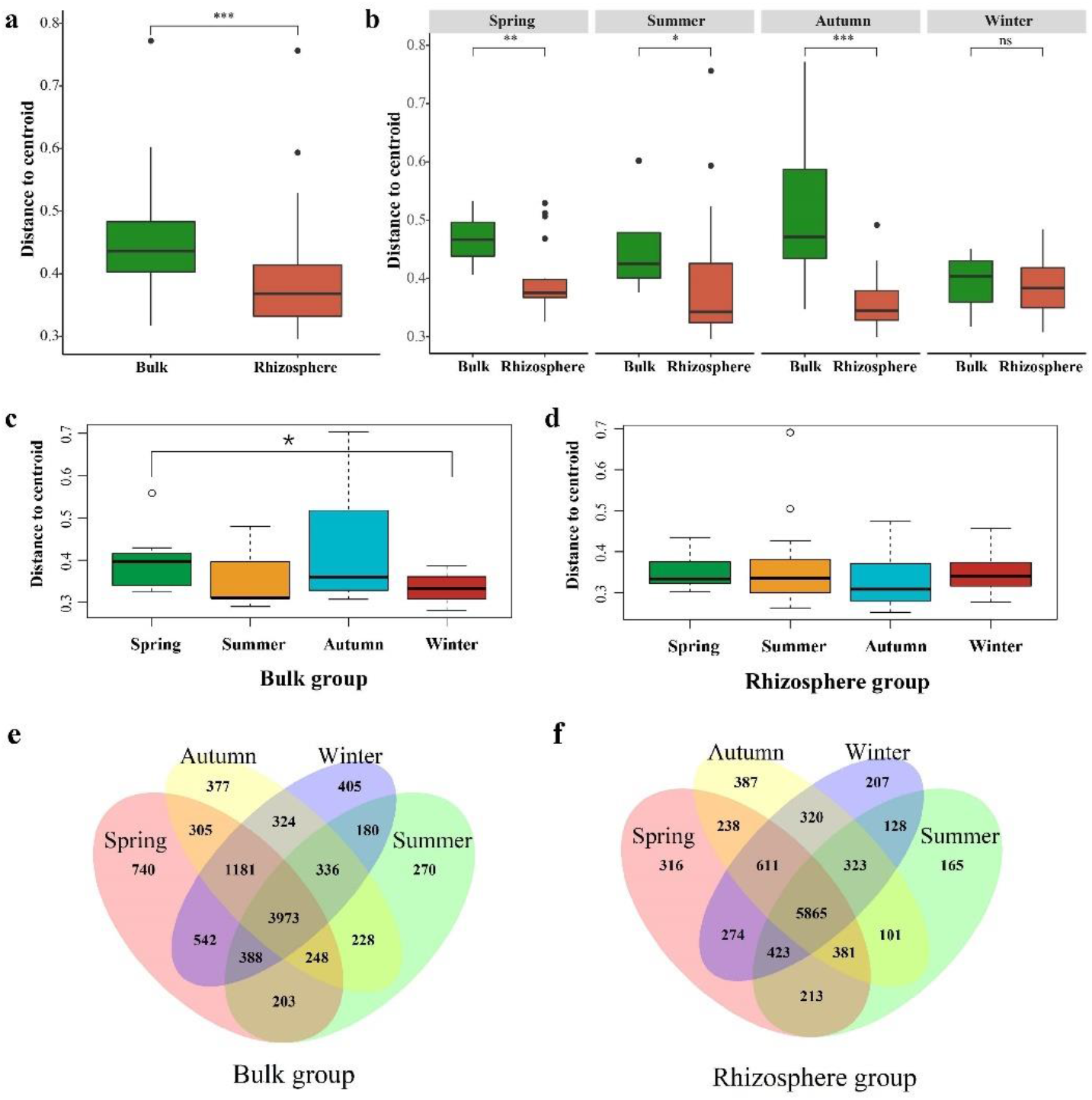
Box plots of Bray-Curtis dissimilarity index. (a-b) The box plot of bacterial community Bray-Curtis dissimilarity indices in bulk and rhizosphere sediment. (c) The box plot of bulk sediment bacterial community Bray-Curtis dissimilarity indices in different seasons. (d) The box plot of rhizosphere sediment bacterial community Bray-Curtis dissimilarity indices in different seasons. Asterisks indicate significant differences between bulk and rhizosphere sediments based on the Wilcoxon rank-sum test. \**P*<0.05; \*\**P*<0.01; \*\*\**P*<0.001. (e-f) Venn Diagram of the number of OTUs of bacterial community in different seasons.

**Figure 3.**
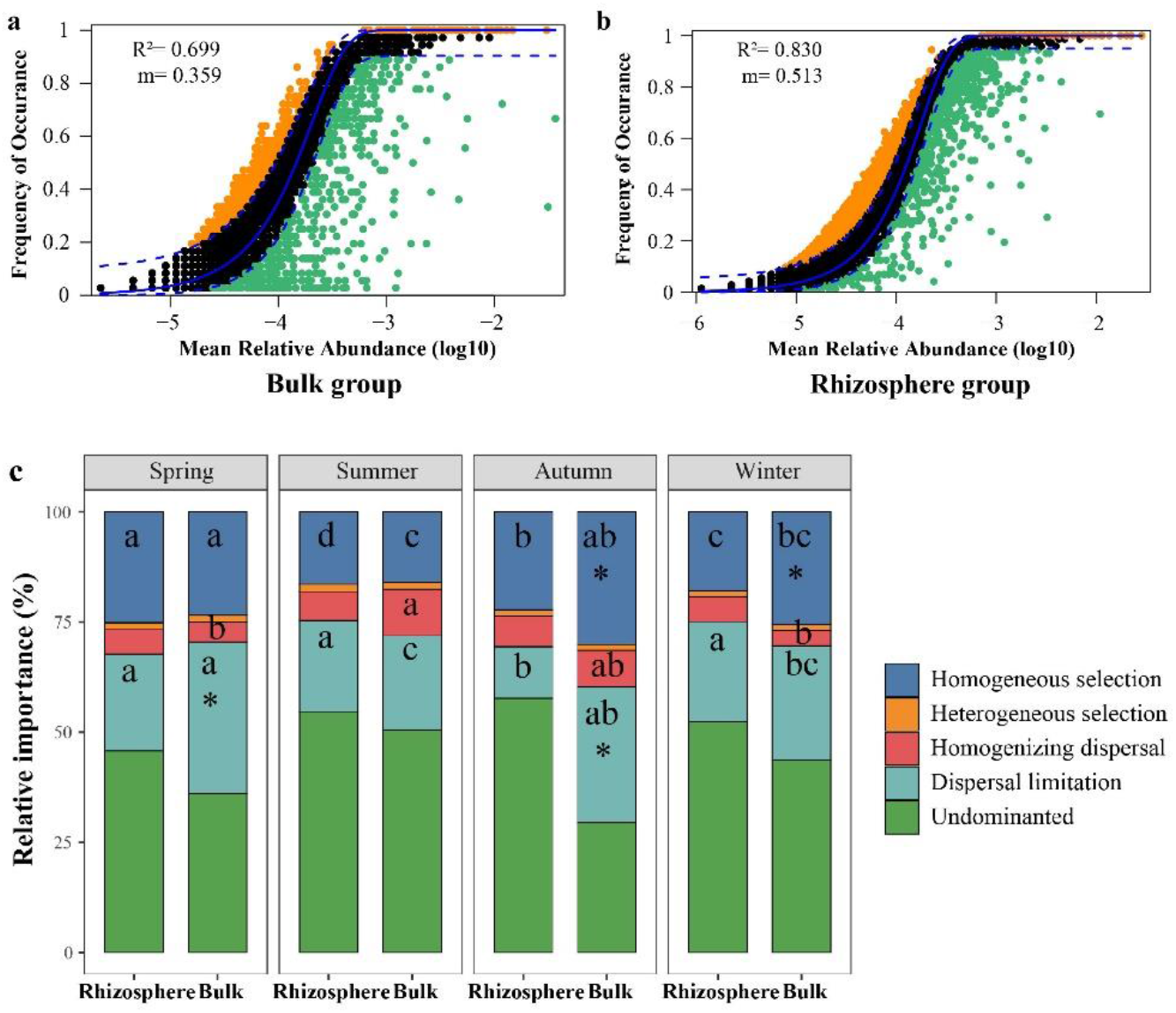
Bacterial community construction. (a-b) Neutral model fitting of OTU in rhizosphere and bulk sediment bacterial communities. (c) Community assembly processes of rhizosphere and bulk bacterial communities in different seasons.

Environmental conditions may have an impact on the assembly of bacterial communities. In terms of the stochastic process, we found that the proportions of homogenizing dispersal increased and dispersal limitation decreased with increasing sediment carbon, and nitrogen concentration. Furthermore, we found that dispersal limitation is positively correlated with environmental distance while homogenizing dispersal is negatively correlated with environmental distance (Figure 4). Hence, higher carbon and nitrogen concentration may lead to greater diffusion capacity. Compared to the bacterial community in rhizosphere sediments, a larger proportion of dispersal limitation and homogenizing dispersal observed in bulk bacterial communities may be due to the lower carbon and nitrogen content in bulk sediments.

**Figure 4.**
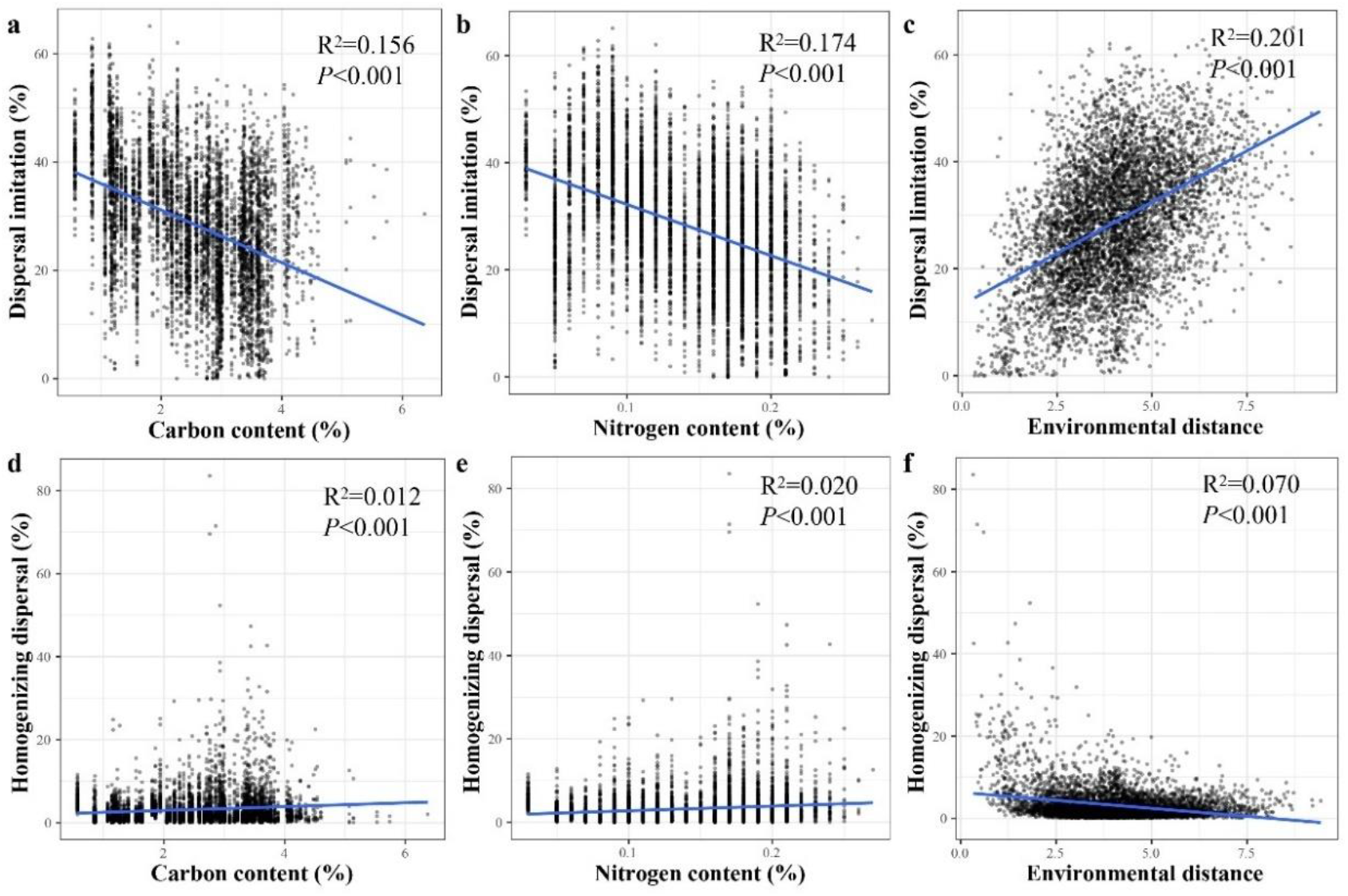
Correlation between bacterial dispersal ability and sediment physicochemical conditions. The correlation between the relative importance of dispersal limitation and homogenizing dispersal with (a, d) carbon, (b, e) nitrogen content and (c, f) environmental distance. For each pair of sediment samples, the lower value of carbon and nitrogen content of the two samples was used.

We found a positive correlation between proportions of dispersal limitation and Bray–Curtis distances and strong negative correlations between proportions of homogenizing dispersal and Bray–Curtis distances (Figure S5 a-b). However, much weaker though significant correlations were observed between proportions of homogeneous selection (or heterogeneous selection) and Bray–Curtis distances (Figure S5 c-d). This implied that dispersal limitation is the major mechanism for increasing the beta diversity of mangrove bacterial communities while homogenizing dispersal is the mechanism for decreasing beta diversity. In mangrove sediments, the carbon and nitrogen content in the rhizosphere sediments was significantly higher than that in the bulk sediments. The higher homogenizing dispersal and lower dispersal limitation caused by high carbon and nitrogen content may lead to the significantly lower beta diversity of the rhizosphere bacterial community.

## 4 Discussion

### 4.1 Influencing factors of bacterial community diversity in bulk and rhizosphere sediments

In this work, we characterized bacterial communities of mangrove sediment samples that covered four seasons across the Dongzhai harbor in Hainan Island. Our results revealed that seasons hold a major role in determining the alpha and beta diversity of sediment bacterial communities. We also found that the alpha diversity in summer was lower than in other seasons. Environmental changes such as temperature, nutrient contents, pH and salinity were important factors affecting the diversity and composition of bacterial communities(Z. Lu et al., 2021; Sunagawa et al., 2015; H. Wang, Gilbert, Zhu, & Yang, 2018). Analyzing the bacterial and fungal communities of global sediment samples, it was found that environmental changes have a strong impact on microbial communities, pH and precipitation were key environmental drivers(Bahram et al., 2018). This study found that the pH was the lowest in summer. Rousk identified the positive relationship between bacterial diversity and higher soil pH, and our results also supported this(Rousk et al., 2010). The seasonal variation of salinity in sediment was mainly related to evaporation and precipitation. Due to the high evaporation caused by the highest temperature, the salinity was the highest in summer. Previous results showed that salinity reduced the diversity of bacterial communities(Ji et al., 2019). In this study, high salinity and low pH in sediment might be the reasons for the low alpha diversity in summer. It can be seen that climate change has an important impact on mangrove bacterial communities.

Although there were obvious separations of the bacterial community composition across different seasons, distinct patterns in bacterial diversity and composition between rhizosphere and bulk sediments were also observed. Plant species were important factors affecting the soil environment. Through litter degradation and root exudation, plants return the nutrients absorbed from the soil to the soil, thus affecting the physicochemical properties of the soil and bacterial community(Fischer et al., 2019; Hobbie, 2015; R. Zhang, Vivanco, & Shen, 2017). Previously research found that mangrove root exudates can increase the content of active organic matter and bacterial activity in the rhizosphere sediment(B. Liu et al., 2017). Besides, tides and waves could also allow mangrove forests to store and transport new fixed carbon(G. et al., 2013; Vo-Luong & Massel, 2008). In this study, significantly higher carbon, nitrogen content and alpha diversity were detected in rhizosphere sediments than that in bulk. The lower organic content and larger fluctuation in bulk sediments could be due to periodic removal of mangrove litter by tidal flooding. It is widely accepted that plant roots, as effective habitat filters, had a strong filtering effect on microbial communities(Davide et al., 2012; B. Liu et al., 2017; Reinhold-Hurek, Bünger, Burbano, Sabale, & Hurek, 2015). As the first barrier of root habitat filtration, the rhizosphere attenuated the impact of environmental pressure on these communities(Davide Bulgarelli, Schlaeppi, Spaepen, Themaat, & Schulze-Lefert, 2013). Our results also confirmed the above conclusion, the beta diversity was significantly lower in the rhizosphere than that in bulk, suggesting convergence in bacterial community composition of the rhizosphere sediments. In terms of alpha and beta diversity, plant roots make the rhizosphere bacterial communities more stable in face of seasonal fluctuation.

### 4.2 Influencing factors of seasonal fluctuations in the relative abundance of rhizosphere and bulk bacteria

Our findings revealed that there were significant differences in the composition of bacterial communities between seasons. There was a clear seasonal fluctuation in the relative abundance of specific bacteria. Some research results indicated that Proteobacteria, Bacteroidetes, and Actinobacteria were significantly enriched in rhizosphere sediments(Palit & Das, 2020; Wu et al., 2016; X. Zhang, Hu, Ren, & Zhang, 2018). It has been confirmed that Actinobacteria and Acidobacteria were dominant in dry season, the relative abundance of Actinobacteria and Acidobacteria increased during the dry period and decreases with the increase of precipitation(Barnard, Osborne, & Firestone, 2013; de Vries et al., 2018). Hainan Island is dry in autumn and winter and rainy in spring and summer, so the relative abundance of Actinobacteria and Acidobacteria was significantly lower in winter than in spring. Our research found that the relative abundance of Verrucomicrobiota was significantly higher in summer compared to the other seasons. Previous studies have found that Verrucomicrobiota was better adapted to high-salt soil environments than other bacteria and that its interaction with salt-tolerant plants plays an important role in mitigating and regulating salt stress(Mukhtar et al., 2018; Szymanska et al., 2018). It has been found that the abundance of Deltaproteobacteria increases with salinity(Behera et al., 2019). Wang showed that the relative abundance of Verrucomicrobiota was negatively correlated with pH, and that Verrucomicrobiota was sensitive to small changes in pH(Martiny, Jones, Lennon, & Martiny, 2015; Rousk et al., 2010). Szymańska studied the composition of the rhizosphere bacterial community under different salinity levels and found that the rhizosphere sediment was enriched with a variety of salt-tolerant bacteria such as Deltaproteobacteria, Calditrichaeota and Fibrobacteres. These bacteria have a potential role in alleviating salt stress in saline plants(Szymanska et al., 2018). The rhizosphere bacterial community mainly contains fast-growing, nutrient-rich bacteria, such as Bacteriodetes, Proteobacteria. The abundance of carbon in the rhizosphere sediment allows them to rapidly growing in plant roots(Sanchez-Canizares, Jorrin, Poole, & Tkacz, 2017). In this study, the relative abundance of Deltaproteobacteria and Calditrichaeota was found to be significantly positively correlated with the carbon content of the sediment and significantly enriched in plant roots.

### 4.3 Bacterial community construction in rhizosphere and bulk sediments

Numerous studies have shown that heterogeneous selection, homogeneous selection, homogeneous dispersal and dispersal limitation dominate the construction of mangrove bacterial communities(C. J. Zhang et al., 2019; Z. F. Zhang, Pan, Pan, et al., 2021; Z. F. Zhang, Pan, Liu, & Li, 2021). We found that dispersal limitation, homogeneous selection, and undominated processes account for a significant proportion of the five ecological processes, dominating the construction of bacterial communities in mangrove sediments. Song also found that homogeneous selection and dispersal limitation play an important role in the construction of bacterial communities in coastal solar salterns(Song et al., 2022). In this study, the undominated process and dispersal limitation dominated the construction of mangrove bacterial communities. These two mechanisms would lead to significant differences in species composition between communities, which also confirms our previous conclusion that species replacement was the reason for the differences in beta diversity. The bacterial communities in mangrove sediment may have adapted to highly dynamic environments disturbed by tides, groundwater, and mud flow. Therefore, community construction may be less affected by environmental factors, resulting in a very low proportion of heterogeneous selection in almost all components(J. Liu et al., 2020). Comparing the construction processes of rhizosphere and bulk bacterial communities, we found that the proportion of homogeneous dispersal of rhizosphere groups was higher and the proportion of dispersal limitation was lower, which is consistent with the results obtained in the NCM. Previous studies have shown that rhizosphere effects may lead to a greater convergence of rhizosphere bacterial community(He, Zeng, Zhao, Wang, & Wu, 2022). In addition, our study found that the carbon and nitrogen content of the sediment was significantly negatively correlated with dispersal limitation and positively correlated with homogeneous dispersal. Several studies have shown that dispersal leads to a reduction in beta diversity of bacterial communities(Catano, Dickson, & Myers, 2017; Mouquet & Loreau, 2003). Our study also showed that homogeneous dispersal was significantly negatively correlated with beta diversity and dispersal limitation was positively correlated with beta diversity. Therefore, the high diffusion ability of rhizosphere bacteria caused by high carbon and nitrogen conditions in sediments, as well as the rhizosphere effects of plants themselves, may be the reason for the significantly lower beta diversity of rhizosphere bacterial communities compared to bulk groups. From the perspective of the influencing factors of community construction, the carbon and nitrogen content in mangrove sediments has an important impact on the construction of bacterial communities.

## 5 Conclusion

This study demonstrates that there are significant differences in the composition and structure of bacterial communities in mangrove sediments between rhizosphere and bulk groups, and both exhibit significant seasonal changes. The assembly of mangrove microbial communities was dominated by stochastic processes. Multiple factors regulate microbial community composition, with pH and carbon, nitrogen content being potentially principal drivers. The high carbon and nitrogen content in rhizosphere sediments may lead to higher homogenizing dispersal and lower dispersal limitation. This may be the reason why the beta diversity of rhizosphere bacterial communities was significantly lower than that of bulk bacterial communities.

## Acknowledgements

We thank Mr. Cairong Zhong, Ms. Ranran Zhu, and Mr. Yulong Li for their help in sampling. This project was supported by the National Natural Science Foundation of China (32170230, 31971540, 31830005, 42276159); the Guangdong Basic and Applied Basic Research Foundation(2023B1515020083); and the Innovation Group Project of Southern Marine Science and Engineering Guangdong Laboratory (Zhuhai) (311021006).

## Supplementary Tables and Figures

**Table S1.** Relative importance of season and sediments contributing to the physicochemical factors by between-subjects effects test on monofactor analysis.

|  | Season |  | Sediment |  |
| --- | --- | --- | --- | --- |
|  | R<sup>2</sup> | P | R<sup>2</sup> | P |
| Salinity/psu | 0.617 | <0.001 | -0.006 | 0.501 |
| pH | 0.163 | <0.001 | 0.032 | 0.068 |
| TC/% | 0.014 | 0.215 | 0.148 | <0.001 |
| TH/% | 0.020 | 0.83 | 0.034 | 0.06 |
| TN/% | -0.016 | 0.722 | 0.193 | <0.001 |
| TS/% | 0.028 | 0.114 | 0.053 | <0.05 |
| Moisture/% | 0.049 | <0.05 | 0.037 | 0.051 |

Note: TC, total carbon content; TH, total hydrogen content; TN, total nitrogen content; TS, total sulfur content.

**Table S2.** Analysis of similarities (ANOSIM) and Permutational multivariate ANOVA (PERMANOVA) results of the bacterial community composition between rhizosphere and bulk sediments based on Bray-Curtis distance, weighted UniFrac distance and unweighted UniFrac distance metrics.

| Distance | Species | ANOSIM |  | PERMANOVA |  |  |  |
| --- | --- | --- | --- | --- | --- | --- | --- |
|  |  | R | P | MeanSqs | F.Model | R<sup>2</sup> | P |
| Bray-Curtis | *A.corniculatum* vs *K.obovata* | 0.02 | - | 0 | 0.3 | 0 | - |
|  | *K.obovata* vs Bulk | 0.2 | *** | 0.11 | 10.3 | 0.13 | *** |
|  | *A.corniculatum* vs Bulk | 0.21 | *** | 0.09 | 7.67 | 0.1 | ** |
| Weighted UniFrac | *A.corniculatum* vs *K.obovata* | 0.02 | - | 0 | 0.29 | 0 | - |
|  | *K.obovata* vs Bulk | 0.12 | *** | 0.07 | 12.97 | 0.16 | *** |
|  | *A.corniculatum* vs Bulk | 0.1 | ** | 0.06 | 10.84 | 0.13 | *** |
| Unweighted UniFrac | *A.corniculatum* vs *K.obovata* | 0.01 | - | 0 | 0.51 | 0.01 | - |
|  | *K.obovata* vs Bulk | 0.11 | *** | 0.01 | 6.81 | 0.09 | *** |
|  | *A.corniculatum* vs Bulk | 0.1 | *** | 0.01 | 5.11 | 0.07 | ** |

**Table S3.**
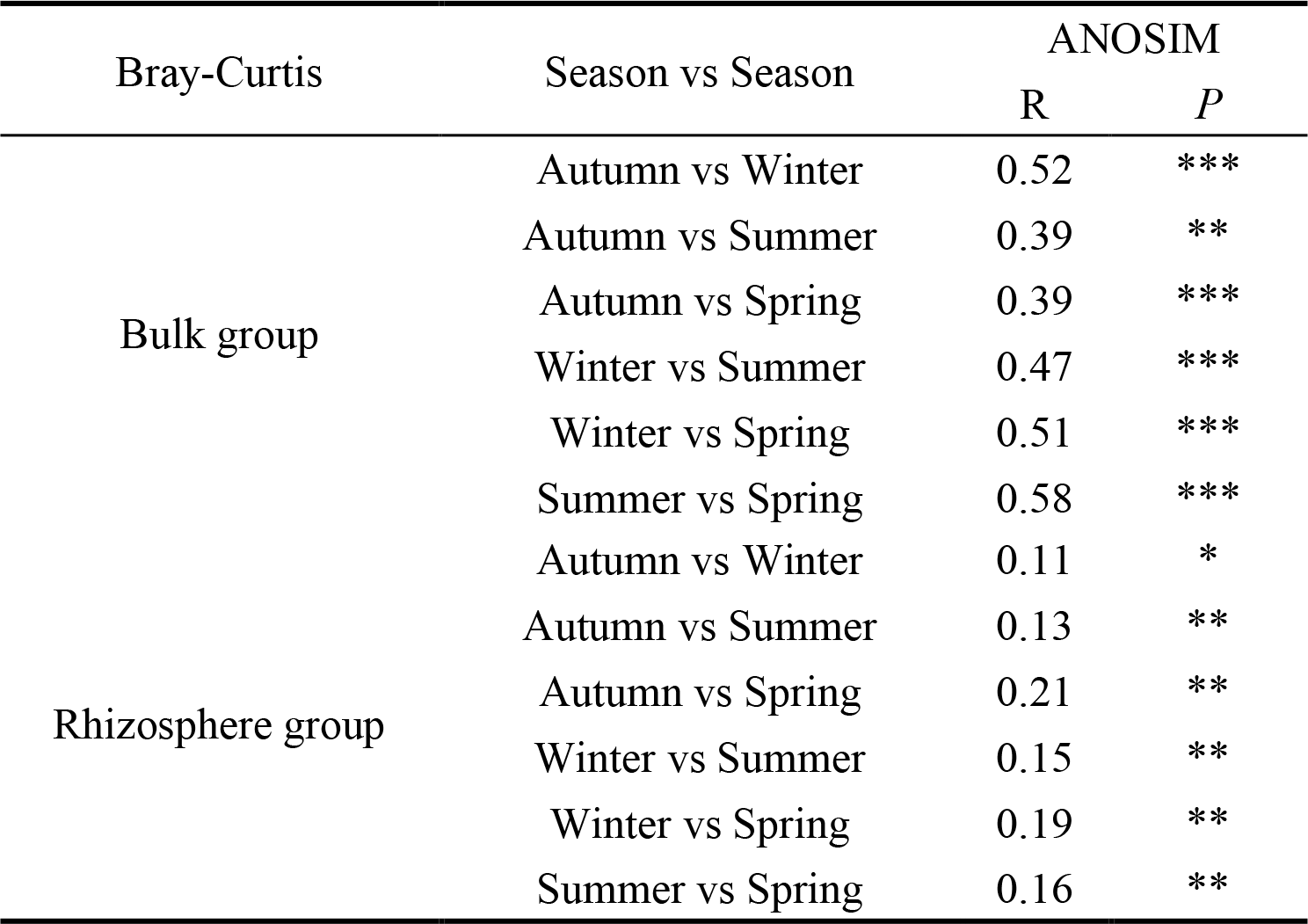
Analysis of similarities (ANOSIM) results of bulk and rhizosphere sediment bacterial community composition among different seasons based on Bray-Curtis distance metrics.

**Table S4.** Analysis of similarities (ANOSIM) results of bulk sediment bacterial community composition among different season based on weighted UniFrac and unweighted UniFrac distance metrics.

| Bulk group | Season vs Season | ANOSIM |  |
| --- | --- | --- | --- |
|  |  | R | P |
| Weighted UniFrac | Autumn vs Winter | 0.44 | *** |
|  | Autumn vs Summer | 0.25 | * |
|  | Autumn vs Spring | 0.49 | *** |
|  | Winter vs Summer | 0.22 | * |
|  | Winter vs Spring | 0.85 | *** |
|  | Summer vs Spring | 0.31 | ** |
| Unweighted UniFrac | Autumn vs Winter | 0.16 | * |
|  | Autumn vs Summer | 0.2 | * |
|  | Autumn vs Spring | 0.2 | * |
|  | Winter vs Summer | 0.59 | *** |
|  | Winter vs Spring | 0.42 | *** |
|  | Summer vs Spring | 0.6 | *** |

**Table S5.**
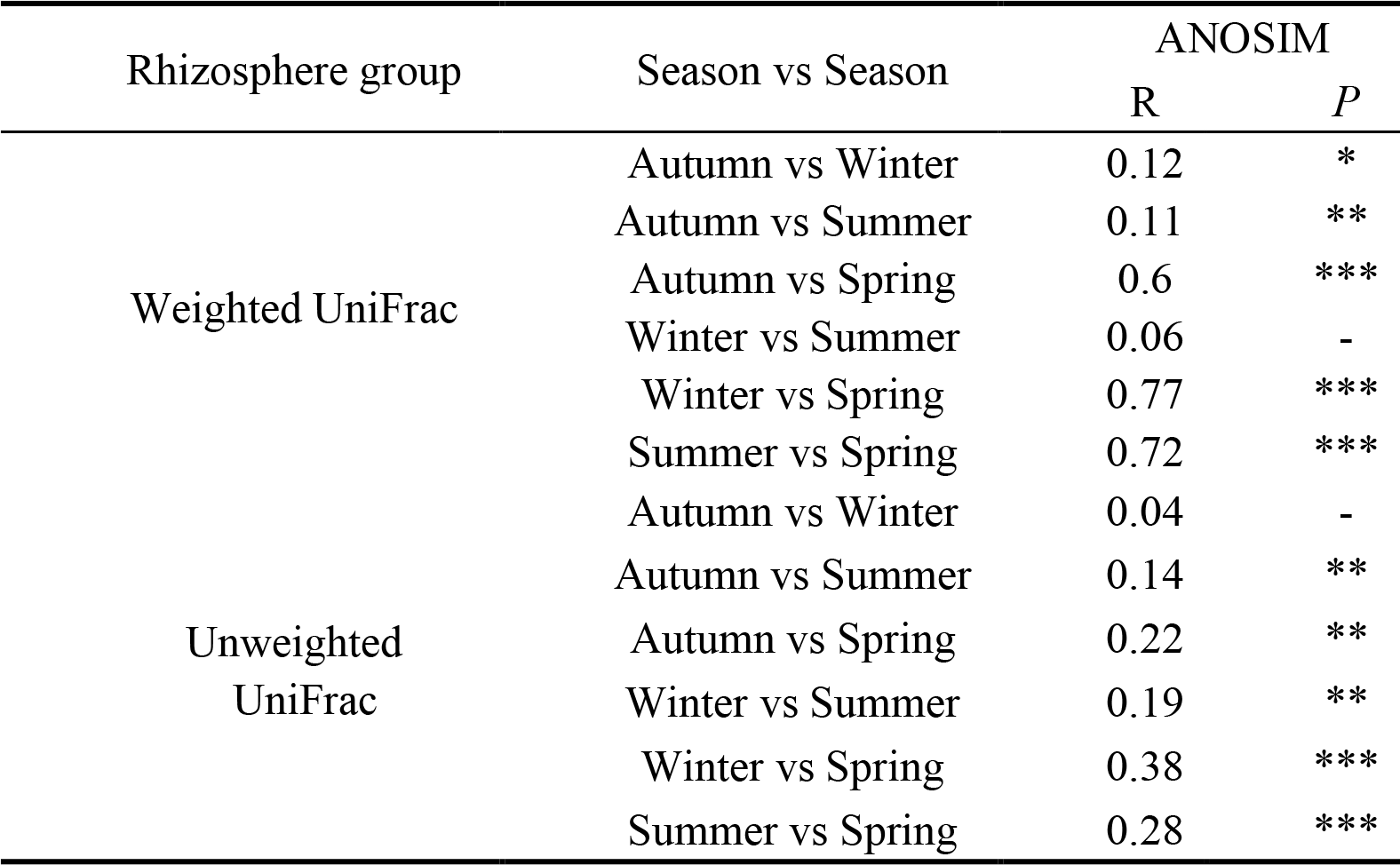
Analysis of similarities (ANOSIM) results based on weighted UniFrac distance and unweighted UniFrac distance of rhizosphere sediment bacterial communities β-diversity among different season.

**Table S6.** Relative importance of stochastic process and deterministic process in rhizosphere and bulk bacterial community construction among different seasons.

| Season | Sediment | Deterministic process (%) | Stochastic process (%) |
| --- | --- | --- | --- |
| Spring | Rhizosphere | 26.57 | 73.43 |
|  | Bulk | 24.99 | 75.01 |
| Summer | Rhizosphere | 18.13 | 81.87 |
|  | Bulk | 17.64 | 82.36 |
| Autumn | Rhizosphere | 23.62 | 76.38 |
|  | Bulk | 31.4 | 68.6 |
| Winter | Rhizosphere | 19.33 | 80.67 |
|  | Bulk | 26.91 | 73.09 |

**Figure S1.**
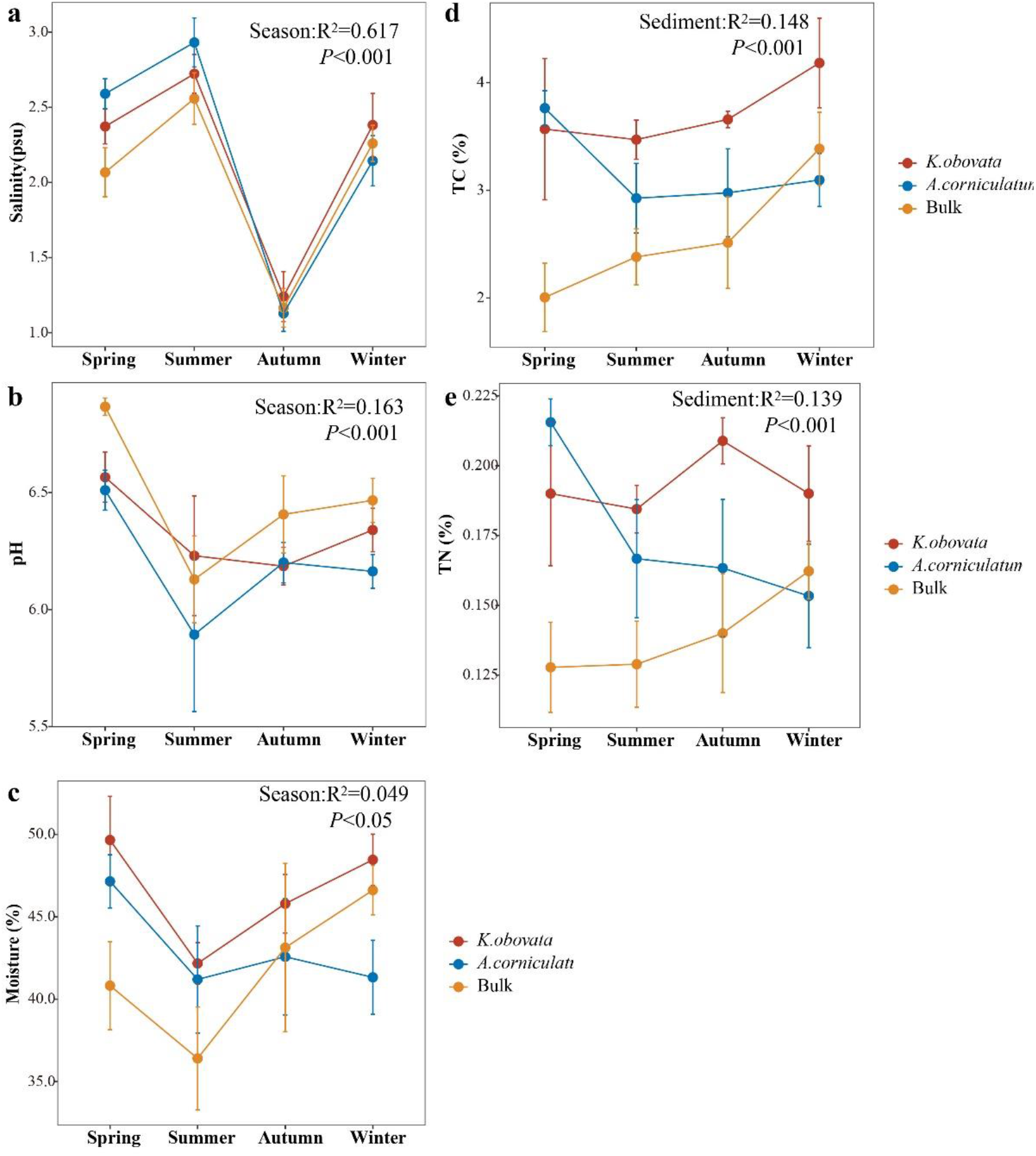
The fluctuation of the mangrove forest sediment physicochemical factors in different season, a is line graphs of the salinity, b is line graphs of the pH, c is line graphs of the moisture, d is line graphs of the total carbon content, e is line graphs of the total nitrogen content. Different colored lines indicate different species.

**Figure S2.**
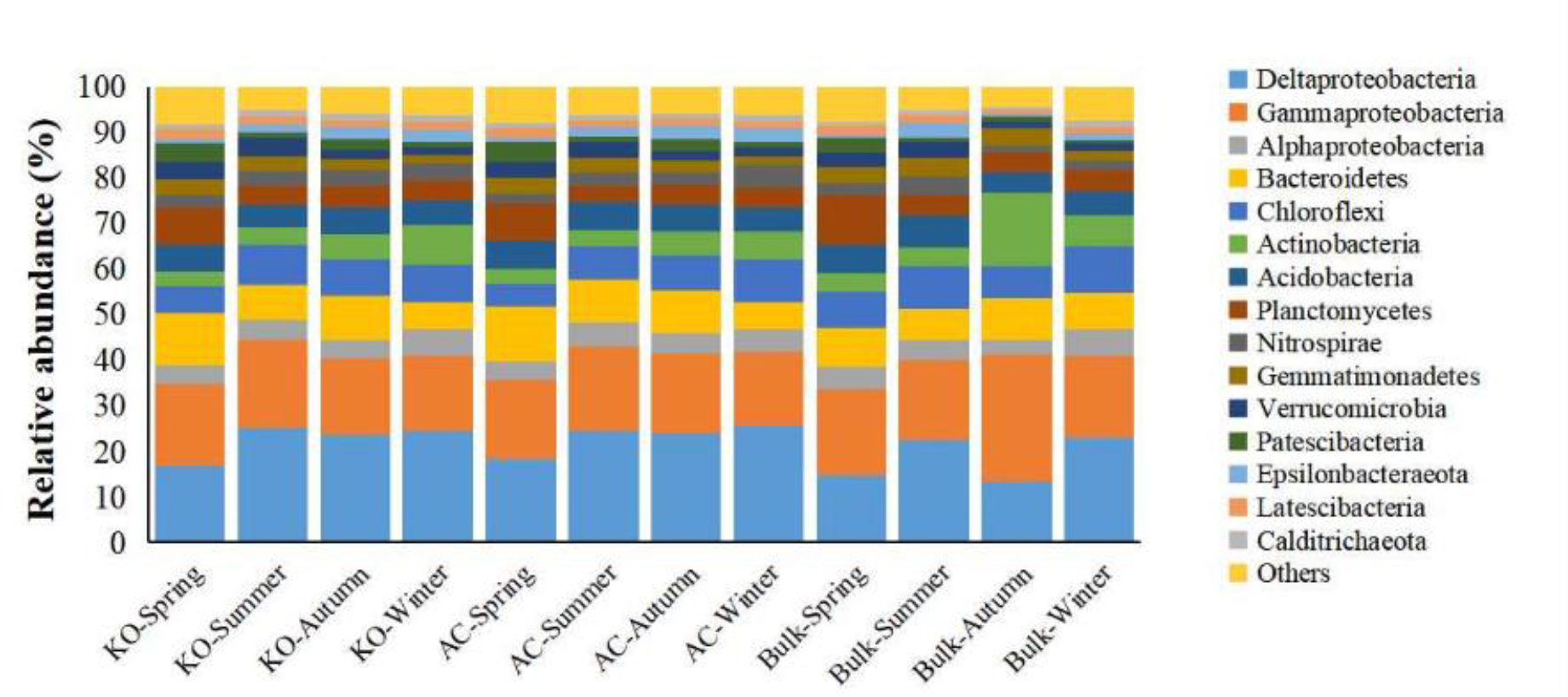
Composition of bacterial communities in the rhizosphere sediments of *Kandelia obovata*, *Aegiceras corniculatum*, and bulk sediments in different seasons.

**Figure S3.**
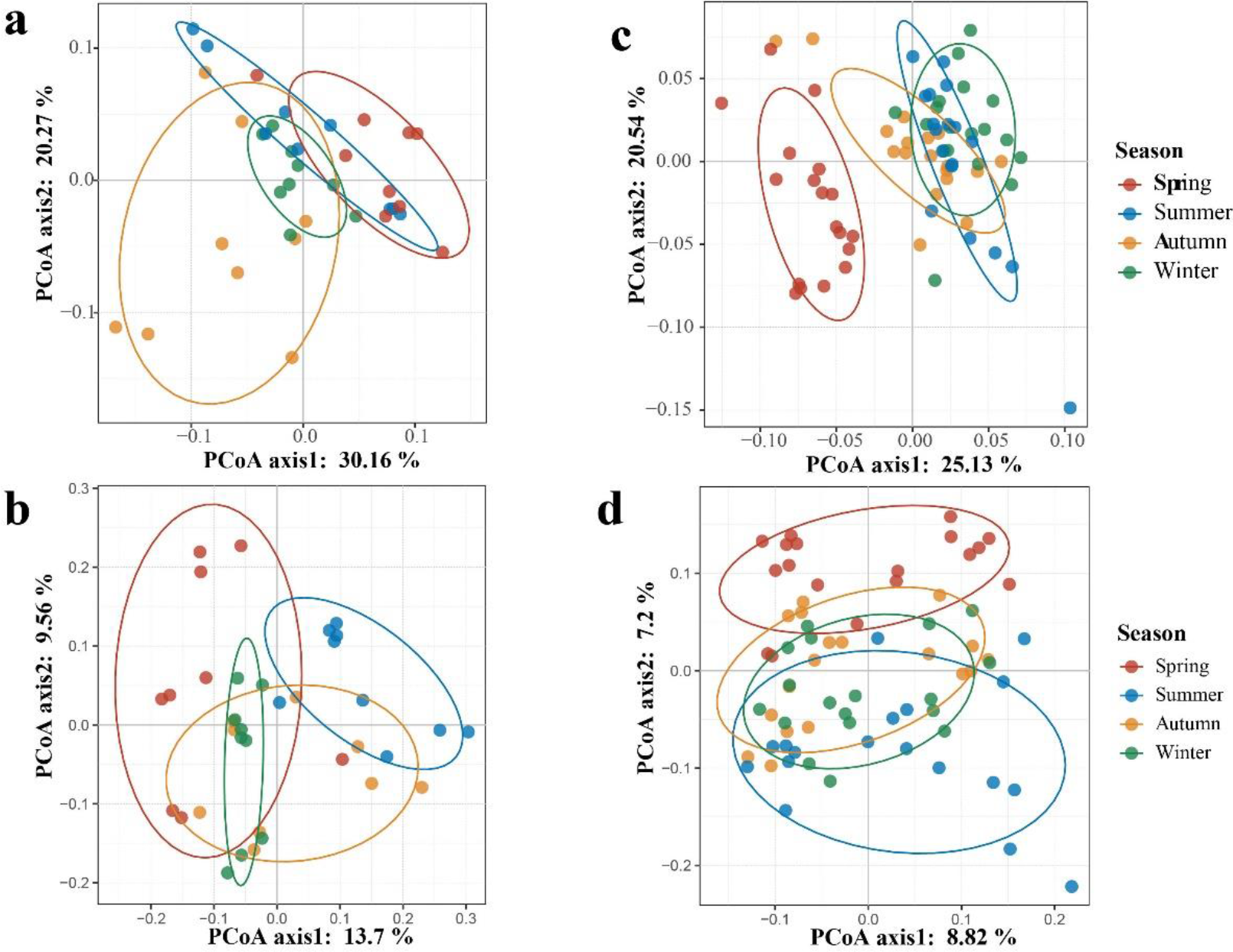
Principal coordinate analysis of bulk and rhizosphere sediment bacterial communities, a is PCoA plots based on Bray-Curtis distance of bulk group b is PCoA plots based on weighted UniFrac distance of bulk group, and c is PCoA plots based on unweighted UniFrac distance of bulk group, d is PCoA plots based on Bray-Curtis distance of rhizosphere group, e is PCoA plots based on weighted UniFrac distance of rhizosphere group, and f is PCoA plots based on unweighted UniFrac distance of rhizosphere group. The X-axis label PCoA axis1 represents the first principal axis that can best distinguish all samples, which can explain the percentage of all differences in the sample; the Y-axis label PCoA axis2 represents the second principal axis that can best distinguish all samples, bacterial communities are color coded according to the season.

**Figure S4.**
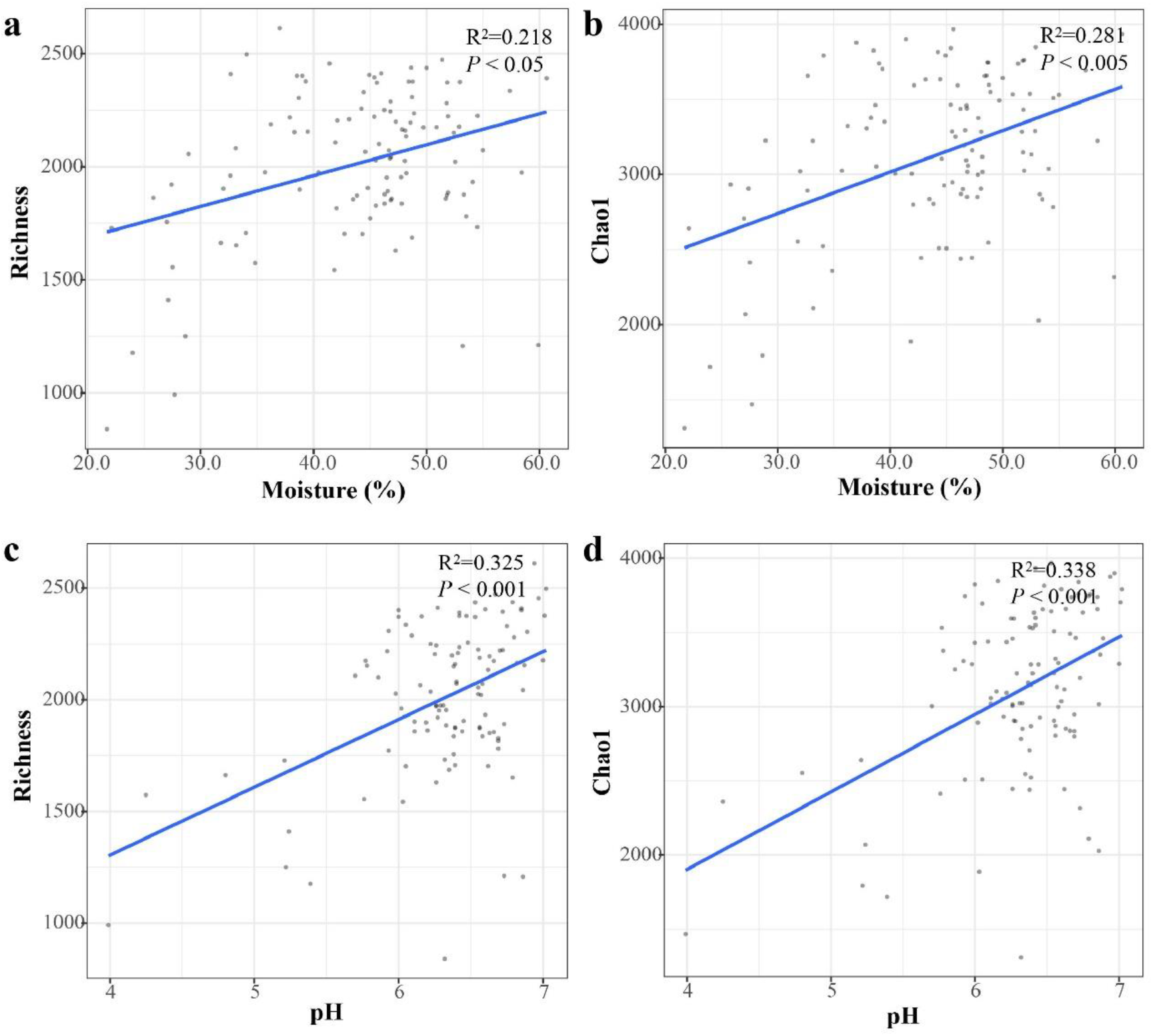
Linear regression analysis diagram of sediment physicochemical properties with bacterial community Chao1 index and Richness. a is linear graph of moisture and richness index, b is linear graph of moisture and Chao1 index, c is linear graph of pH and richness index, d is linear graph of pH and Chao1 index.

**Figure S5.**
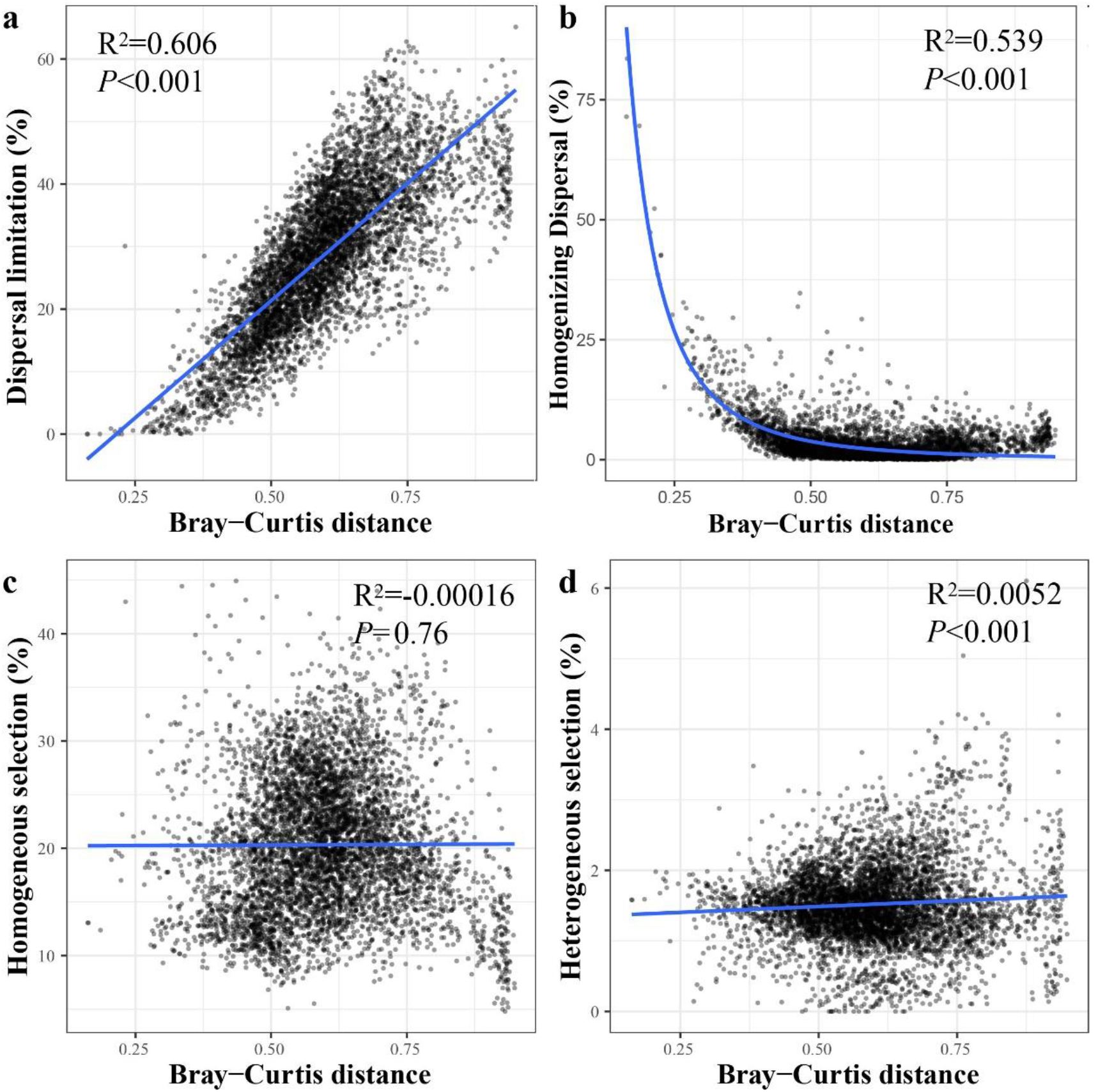
The correlation between the relative importance of dispersal limitation (a), homogenizing dispersal (b), homogeneous selection (c) and heterogeneous selection (d) with Bray-Curtis distance.

